# Gating effects of a Cav2.3 calcium channel mutation linked to developmental and epileptic encephalopathy

**DOI:** 10.64898/2026.05.25.727692

**Authors:** Devon Khousakoun, Ivana A. Souza, Laurent Ferron, Maria A. Gandini, Gerald W Zamponi

## Abstract

Developmental and epileptic encephalopathies (DEEs) are a group of neurological disorders primarily affecting young children and are characterized by severe seizures. DEEs are challenging to manage, with some patients experiencing severe side effects or not responding to frontline therapies. This is partly because of the many underlying mechanisms involved in DEE pathology and the relatively limited mechanism-specific action of current treatments. The *CACNA1E* gene, which encodes the voltage-gated calcium channel Cav2.3 (R-type), has recently been associated with DEEs. More than fifteen different mutations in *CACNA1E* have been identified in patients with DEEs; however, the mechanisms by which these mutations affect channel function and, thus, their relationship to DEEs, remain largely unknown. Previous research has begun to characterize the functional effects of R-type channel mutations on channel biophysics, but only a handful of mutations have been studied functionally to date. Here, we transiently expressed Cav2.3 channels and used whole-cell patch-clamp to examine the biophysics of one specific disease-associated R-type channel mutant in which leucine 228 is substituted with a proline (L228P). Compared to wild-type, the L228P mutant did not alter peak current density, inactivation kinetics, or recovery from inactivation, but showed a significant shift towards hyperpolarized voltages in both voltage-dependent activation and steady-state inactivation. This resulted in a broader window current shifted towards more hyperpolarized potentials, which predicts increased channel availability and activity at subthreshold voltages relative to wild-type channels. Our results contribute to the ongoing characterization of R-type mutants, with the long-term goal of informing mechanism-specific therapies for DEEs.

## Introduction

Developmental and epileptic encephalopathies (DEEs) are a group of debilitating neurological disorders that are typically characterized by seizures, delayed or impaired cognitive development, and abundant epileptiform activity on electroencephalogram (EEG) (McTague et al., 2016; Raga et al., 2021). These disorders disproportionately affect young children, with high mortality rates and a significant impact on quality of life. Despite the profound impact of these conditions, treatment options remain limited, in part, due to their genetic heterogeneity (Guerrini et al., 2023). Most cases arise from *de novo* mutations in a wide range of genes, resulting in diverse underlying mechanisms. Given this, there is a need for precision therapies that address the specific underlying deficits. This is where current treatments fall short. Front-line treatment includes anti-seizure medications (ASMs), many of which target voltage-gated calcium or sodium channels. However, these treatments are often non-specific, failing to address the distinct functional consequences of pathogenic variants. As a result, some patients do not respond at all, and others develop severe side effects or strong tolerance over time (Samanta et al., 2025).

Mutations in the *CACNA1E* gene, which encodes the Cav2.3 (R-type) voltage-gated calcium channel, have been identified in patients with DEE conditions (Helbig et al., 2018). Initial functional studies of four Cav2.3 mutants revealed altered activation gating and channel kinetics (Helbig et al., 2018) that were largely consistent with gain-of-function. On the other hand, a recent study that investigated three additional variants reported loss-of-function effects (Souza et al., 2026). To further understand how Cav2.3 channel mutations affect pathogenicity, it is thus important to characterize additional variants. Cav2.3 channels have been implicated in neuronal excitability, specifically in brain areas such as CA1 hippocampal neurons, where they play a critical role in action potential burst generation (Magee and Carruth, 1999). Indeed, upregulation of intrinsic calcium-dependent bursting in CA1 hippocampal neurons may contribute to hippocampal epileptogenicity (Sanabria et al., 2001). Cav2.3 channels have also been associated with synaptic plasticity (Breustedt et al., 2003; Dietrich et al., 2003). Cav2.3-deficient mice exhibited significant decreases in oscillatory burst discharges in reticular thalamic neurons, thereby limiting the expression of an absence epilepsy phenotype (Zaman et al., 2011). Multiple studies have also shown that Cav2.3-deficient mice exhibit marked resistance to seizures (Weiergräber et al., 2006, 2007), while otherwise showing only a mild phenotype including altered sleep patterns, increased timidness, and hyposensitivity to pain (Saegusa et al., 2000; Siwek et al., 2014).While there is growing evidence showing the role of Cav2.3 in epilepsy-related processes, it is still unknown how disease-associated mutations alter channel functions and contribute to DEE pathophysiology.

The present study aimed to characterize the functional impact of the L228P *CACNA1E* mutation located within the S5 segment of domain I (Helbig et al., 2018) (**Figure 1a**), a region expected to contribute to channel gating. Using whole-cell patch-clamp electrophysiology in a tsA-201 cell expression system, L228P mutant channel biophysics were compared with wild-type Cav2.3 channels. We show that the L228P mutation alters channel activity and availability across a broad range of membrane voltages compared to wild-type channels by inducing a ∼30 mV hyperpolarizing shift in half-activation and inactivation potentials. Our results contribute to the ongoing functional characterization of *CACNA1E* mutations and the long-term goal of developing precision therapies to treat DEEs.

**Figure 1.**
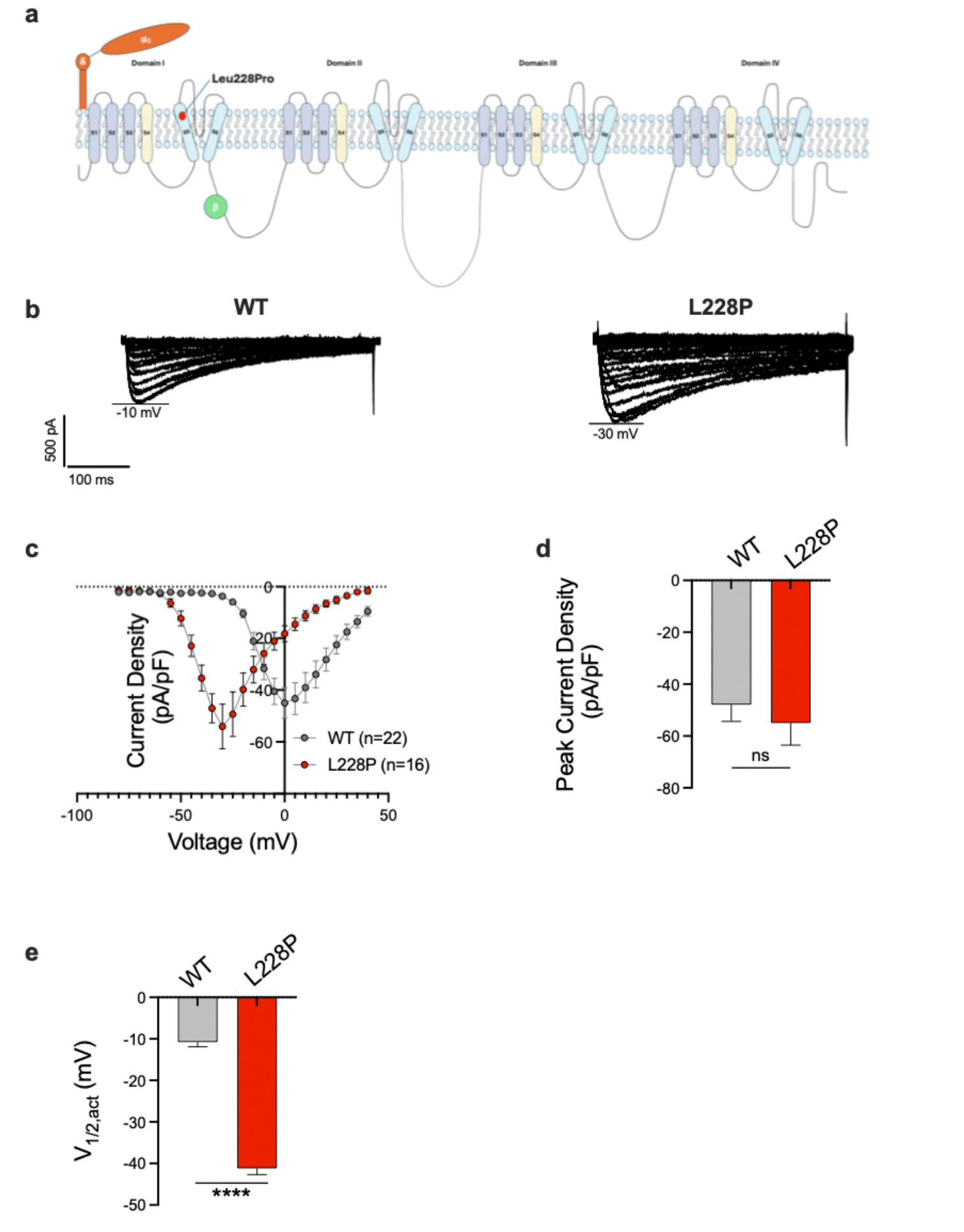
a) Cav2.3 α1 subunit membrane topology, with the location of the L228P mutation. b) Families of representative whole-cell current traces for a cell expressing wild-type Cav2.3 channels (left) and L228P mutant Cav2.3 channels (right). The horizontal bars indicate the voltage step at which peak inward current was observed for each condition. c) The current-voltage (I-V) relationship showing mean current density as a function of voltage for wild-type and L228P mutant channels. The same cells were used for the analyses shown in panels d) and e). d) A comparison of the mean peak current density between wild-type and L228P mutant channels. e) A comparison of the half-activation voltage (V_1/2,act_) between both conditions. Data are presented as mean ± SEM. Statistical significance is indicated as **** (*p*<0.0001); ns = not significant.

## Methods

### cDNA Transfection

Human embryonic kidney tsA-201 cells were grown in Dulbecco’s Modified Eagle’s Medium (DMEM), containing 10% fetal bovine serum and 1% penicillin/streptomycin. The cells were maintained at 37ºC and in a humidified atmosphere containing 5% carbon dioxide. Cells were suspended in a medium containing 0.25% trypsin and ethylenediaminetetraacetic acid (EDTA) and plated on glass coverslips in 10-centimetre culture dishes 6 hours prior to transfection. The calcium-phosphate method was utilized to transfect cells (Kwon and Firestein, 2013). In all conditions, cells were transfected using 3 µg each of cDNA plasmids encoding the Cavβ1b subunit and Cavα2-δ1 subunit, along with 3 µg of Cav2.3α1 subunit cDNA (wild-type or mutant). Furthermore, all conditions were co-transfected with 0.5 µg of green fluorescent protein (GFP) cDNA to facilitate identification of transfected cells. The day after transfection, the cells were moved to 30ºC and kept in a humidified atmosphere containing 5% carbon dioxide. Cells were utilized 72 hours post-transfection and then discarded. Cell culture and cDNA transfection methods were adapted from Harding et al. (2023).

To engineer the mutants into the human Cav2.3 channel (α1E-3 splice variant (GenBank: [http://ncbi-n:NM_001205293.1]), site-directed mutagenesis was performed by GenScript (Piscataway, NJ, USA). Furthermore, the plasmid was sequenced to verify the presence of the desired variant and exclude any additional mutations.

### Electrophysiology Recordings

Whole-cell patch clamp recordings were conducted at room temperature (22-24ºC), and currents were measured using an Axopatch 200B amplifier linked to a computer containing pCLAMP9.2 software. Borosilicate glass patch pipettes were pulled and polished using an electronic micropipette puller. For patch clamp recordings, only pipettes with a resistance reading of 2.5-5 MΩ were used to ensure good access resistance. Cover slips containing the tsA-201 cells were transferred to circular petri dishes (35 mm x 10 mm) and bathed in external solution. The external solution contained (in mM): 2 CaCl_2_, 1 MgCl_2_, 10 4-(2-hydroxyethyl)-1-piperazineethanesulfonic acid (HEPES), 10 glucose, and 137 CsCl. Recording pipettes were filled with internal solution containing (in mM): 130 CsCl, 2.5 MgCl_2_, 5 EGTA, 10 HEPES, 3 ATP, and 0.5 GTP. When conducting recordings to test for calcium-dependent inactivation (CDI), the same internal solution was used, but with 0.5 mM EGTA. Both the external and internal solutions were adjusted to a pH of 7.4 to maintain physiological conditions. Upon formation of a high-resistance seal (>1 GΩ), the cell membrane was ruptured by suction, and typically only cells with a leak current of less than -200 pA at -100 mV were used for complete recordings. Before recording, cells were allowed to dialyze for one minute to allow for equilibration of the pipette internal solution with the cytoplasmic environment. Series resistance was compensated to 85%, and any cell with a voltage error greater than 5 mV was discarded. To acquire current-voltage (I-V) relations, a step voltage protocol was applied. The cell was held at a holding potential of -100 mV for 200 ms, followed by voltage steps from -80 mV to +40 mV in +5 mV increments. Each voltage step consisted of a 400-ms pulse, followed by a return to the holding potential of -100 mV for 150 ms. Sweeps were delivered at a frequency of 1 Hz. To examine the voltage dependence of inactivation, steady-state inactivation (SSI) curves were acquired using a voltage paradigm consisting of a series of conditioning prepulses, followed by a test pulse. In the SSI protocol for wild-type channels, cells were held at a holding voltage of -100 mV. Then, 5-s conditioning prepulses were applied in 5 mV increments from -110 mV to -15 mV, each followed by a 140-ms test pulse to 5 mV. In the SSI protocol for the L228P mutant, cells were held at a holding voltage of -100 mV, followed by a series of 5-s conditioning prepulses. These prepulses were applied in 5 mV increments from -120 mV to -15 mV, each followed by a 140-ms test pulse to -30 mV. The test pulse was set to the voltage corresponding to the average peak current density on each channel’s I-V curve. A paired-pulse recovery protocol was used to measure recovery from inactivation (RFI). A 2000-ms test pulse to 5 mV was applied, followed by a return to the holding potential of -100 mV. The recovery interval varied, with times (in ms): 10, 20, 40, 80, 160, 320, 640, 1280, 2560, 5120, 10240, and 20480. After each recovery interval, a 150-ms test pulse of the same voltage as the initial test pulse was applied, followed by a return to the holding potential of -100 mV for 100-ms before the protocol repeats with the next recovery interval.

### Data Analysis

Current-voltage values were copied from pCLAMP9.2 software into a Microsoft Excel spreadsheet. Current values for each corresponding voltage were normalized by dividing by the cell’s capacitance. These values were then imported into GraphPad Prism, where current-voltage relationships were plotted and fitted using a Boltzmann equation. The I-V curves were fitted with the following Boltzmann equation: 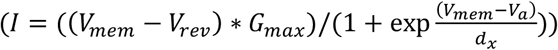 where *I* is the peak current, V_mem_ is the membrane potential, V_rev_ is the reversal potential, G_max_ is the maximum conductance, V_a_ is the half-activation voltage, and d_x_ is the slope factor. When fitting steady-state inactivation curves, the following Boltzmann equation was used: 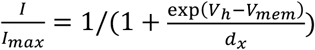, where V_h_ is the half-inactivation voltage. When analyzing RFI, the ratio of the peak of the second test pulse to the peak of the first test pulse was graphed against time and fitted with the following monoexponential equation: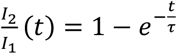, where *I*_0_ is the peak of the second test pulse, *I*_1_ is the peak of the first test pulse, *t* is the recovery interval, and τ is the time constant of RFI. To analyze inactivation kinetics, the decay phase of the current during each depolarizing voltage step was fitted with the monoexponential function: 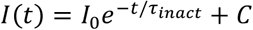, where *I*(*t*) is the current amplitude at a given time after the depolarizing step, *t* is the time after onset of the depolarizing pulse, *I*_2_ is the initial amplitude of the inactivation portion of the current, τ_inact_ is the time constant of inactivation, and *C* is the non-inactivating portion of the current.

### Statistical Analysis

GraphPad Prism software (version 10.4.1) was used for all statistical analyses. All values were used in the analysis unless they were considered outliers, which was tested via Grubb’s Test for Outliers (α=0.05). Data were assessed for normality using the Shapiro-Wilk test and by visual inspection of Q-Q plots. Data were considered to be approximately normally distributed. Data are expressed as mean ± SEM. *n* represents individual cells. A significance threshold of *p*<0.05 was used. When comparing data between groups, an unpaired two-tailed Student’s t-test was used to test for significant differences.

## Results

### The L228P shifts Cav2.3 activation and steady-state inactivation towards hyperpolarized voltages

Both wild-type and mutant Cav2.3 channels yielded robust whole cell currents (**Figure 1b**); however, peak current for the mutant was elicited at more hyperpolarized potentials. This is also evident upon inspection of the current-voltage (I-V) relationships for wild-type and L228P mutant Cav2.3 channels (**Figure 1c**), where a robust leftward shift in the activation range of the channels is clearly evident. This effect produces a massive gain-of-function effect at hyperpolarized potentials compared to wild-type channels (at least when cells are held at -100 mV). However, despite the much larger driving force at the peak of the L228P I-V curve, peak current density was similar between wild-type and mutant channels (*p*=0.5177, **Figure 1d**), and thus at more depolarized potentials, the mutant channels exhibit a reduced whole-cell current amplitude compared to the wild-type channel at a given voltage. The I-V relation of the mutant channel exhibited a somewhat unusual biphasic decline at more depolarized voltages, with an initial steep phase that extrapolated to around 0 mV, and then a more gradual decline towards the reversal potential for calcium. The initial steep phase may potentially be due to an increased permeability to the cesium ions in the recording solution, a possibility that we did not explicitly explore further here. As a result, we were unable to compare G_max_ values between the mutant and the wild-type channels. Be that as it may, at more positive potentials, the mutant channel appeared to undergo a loss of function relative to wild-type channels. Analysis of the half-activation voltages revealed a significant hyperpolarizing shift (∼30 mV) in the half-voltage of activation (V_1/2,act_) of the mutant compared to wild-type channels (*p*<0.0001, **Figure 1e**).

Steady-state inactivation curves of the wild-type and L228P mutant channels (**Figure 2a**) exhibited no difference in slope factor (d_x_) (*p=*0.6447, **Figure 2b**). However, L228P mutant channels exhibited a significant hyperpolarizing shift of approximately 30 mV in the half-voltage of inactivation (V_1/2,inact_) compared to wild-type channels (*p<*0.0001, **Figure 2c**). This leftward shift in the voltage-dependence of inactivation is consistent with a loss-of-function, such that most channels would be tonically inactivated near typical neuronal resting membrane potentials. The mutant L228P channel also appeared to exhibit a small fraction of non-inactivating current, which was not observed in wild-type channels, and would constitute a gain-of-function.

**Figure 2.**
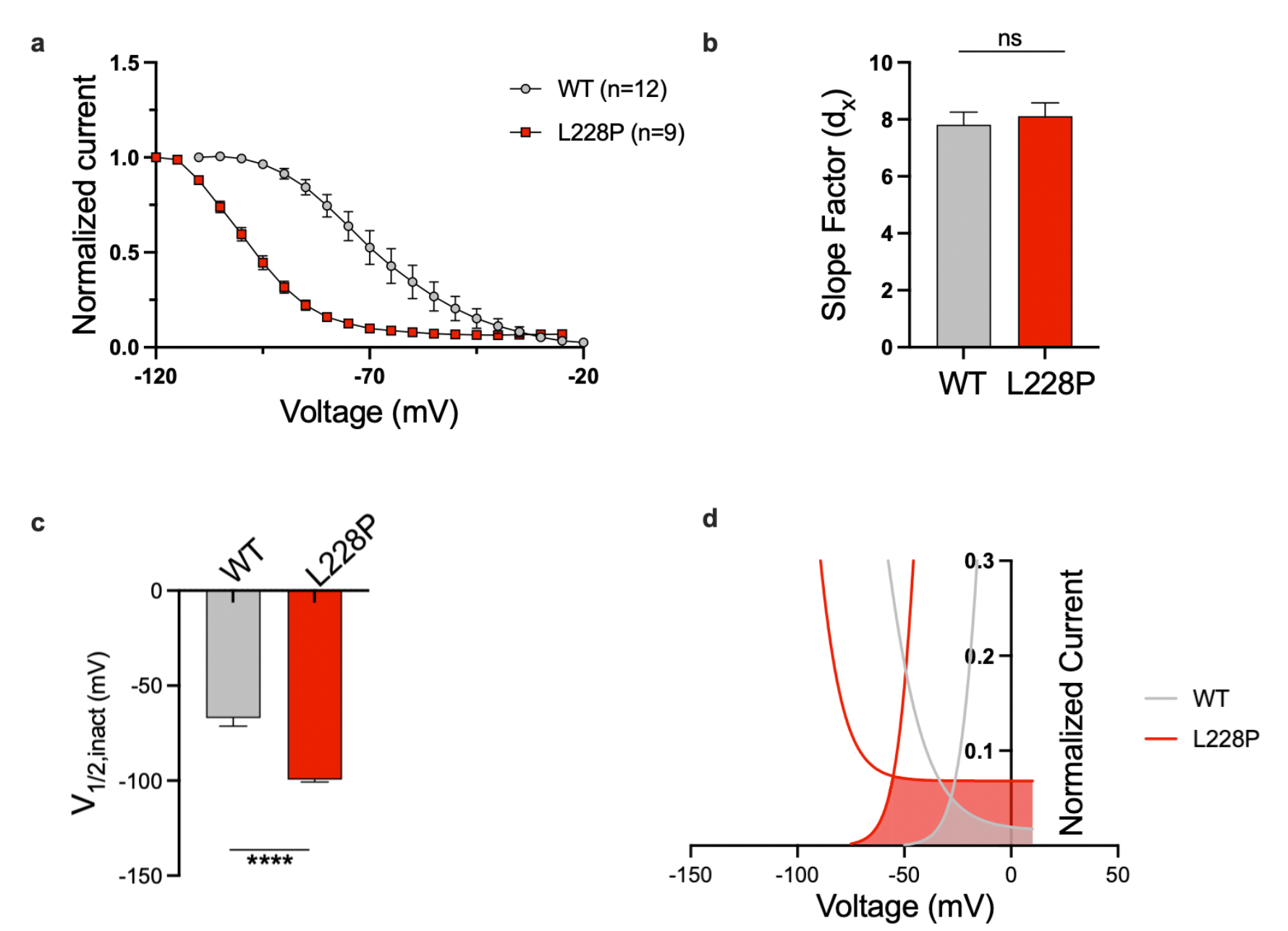
a) Steady-state inactivation curves fitted with a Boltzmann equation for wild-type and L228P mutant channels. b) A comparison of the slope factor (d_x_) between wild-type and L228P channels. c) A comparison of the half-inactivation voltage (V_1/2,inact_) between wild-type and L228P channels. d) Window current plots for wild-type and L228P channels. Data are presented as mean ± SEM. ns = not significant. Statistical significance is indicated by **** (*p*< 0.0001).

To further explore the activation and inactivation properties of the L228P mutant, the window currents of both the L228P mutant and wild-type Cav2.3 channels were plotted (**Figure 2d**). The average area under the curve for the L228P mutant was approximately 290% greater than that of the wild-type current and shifted towards hyperpolarized potentials compared to wild-type channels. As noted above, the SSI curve revealed a non-inactivating fraction of channels across a broad voltage range, which resulted in a broadening of the window current plot.

### The L228P mutation does not alter inactivation kinetics or recovery from inactivation

Because the L228P mutant and wild-type channels activate and inactivate at significantly different voltage ranges (**Figure 1**), the inactivation kinetics could not be directly compared across the same set of voltages. Nonetheless, the time constants for inactivation for the mutant and wild-type channels appeared to be similar, with time constants of around 200 ms. Previous work from the Yue laboratory (Liang et al., 2003) has revealed that rat Cav2.3 channels can undergo calcium-dependent inactivation, but this is only observed when intracellular calcium buffering capability is low. To test whether the mutant affected calcium-dependent inactivation, we thus strived to compare inactivation kinetics for each channel in solutions containing either 5 mM or 0.5 mM EGTA in the recording pipette. For cells expressing wild-type channels, the voltage steps ranged from -10 mV to +10 mV. Under reduced intracellular calcium buffering, wild-type Cav2.3 channels showed a more rapid rate of inactivation at +10 mV only (*p*=0.0305), with no differences in the rate of inactivation being observed at the other voltages tested (all *p*>0.05) (**Figure 3a**). This was surprising and indicates that the human Cav2.3 isoform used in our study does not exhibit robust CDI. For the L228P mutant, the decay phase of voltage steps ranging from -25 mV to -5 mV was used, and no significant differences in the time constant of inactivation were observed at any voltages (*p*>0.05), irrespective of intracellular calcium buffering capacity (**Figure 3b**). Similarly, RFI did not differ between wild-type and L228P mutant channels (**Figure 3c**), with no differences in the time constant of recovery (*p*=0.1237, **Figure 3d**).

**Figure 3.**
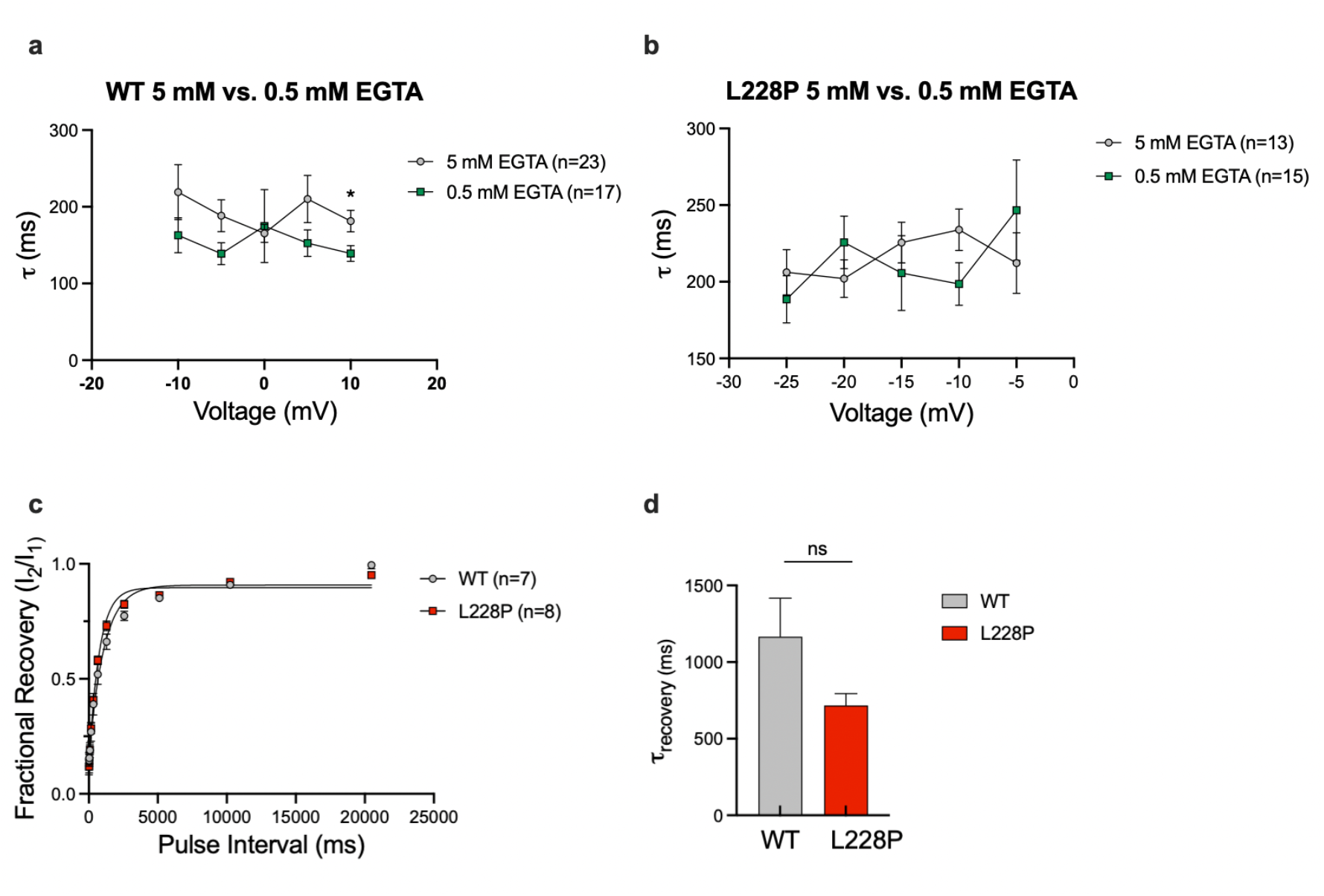
a) Time constant of inactivation plotted as a function of voltage in 5 mM EGTA and 0.5 mM EGTA for wild-type Cav2.3 channels. b) Time constant of inactivation plotted as a function of voltage in 5 mM EGTA and 0.5 mM EGTA for L228P mutant Cav2.3 channels. c) Mean RFI for wild-type and L228P mutant Cav2.3 channels, shown as fractional recovery versus test pulse interval. d) Time constant of recovery for wild-type and L228P mutant Cav2.3 channels. Data are presented as mean ± SEM. Statistical significance is indicated as * (*p*<0.05). ns = not significant.

## Discussion

Here, we report complex gating behaviour of a pathogenic Cav2.3 mutant linked to DEEs. This variant was originally identified in a three-year-old male patient with myoclonic seizures and focal impaired awareness seizures that first occurred at two weeks. EEG showed multifocal discharges, and white matter volume loss was observed by MRI. The patient exhibited spastic quadriplegia and hypotonia, along with profound neurodevelopmental disorder. The patient was nonverbal and nonambulatory, but there was no movement disorder, and there were no congenital contractures (Helbig et al., 2018). Clearly, this Cav2.3 variant triggered severe neuropathology that hinted at a high likelihood of there being major functional changes in the properties of this channel. Our work here confirms this hypothesis, revealing massive shifts in the voltage-dependences of activation and inactivation, along with substantial alterations in the size and position of window current that are largely consistent with a mixed gain- and loss-of-function phenotype.

Peak current density was not different between wild-type and L228P mutant channels. However, the L228P mutant channel activates at much more hyperpolarized voltages, with the peak current density occurring at -30 mV compared to 0 mV for the wild-type channel. At the peak current density for L228P, a much greater electrochemical driving force is expected, and thus, a larger peak current density would also be expected, but this was not observed. In principle, this could be due to an overall reduction in either surface channel expression, open probability, or single-channel conductance. However, the I-V relations were obtained by holding cells at -100 mV, a voltage at which, according to the steady-state inactivation data in **Figure 2**, only approximately 60 percent of the channels are available for opening. Indeed, this reduced availability can fully account for the observation that peak current densities of wild-type and mutant channels were similar despite the difference in driving force (i.e., had all channels been available for opening, peak current density of the mutant channels at -30 mV would be expected at ∼ -85 pA/pF). As noted earlier, G_max_ could not be derived for the L228P mutant because it assumes a linear relationship between current and voltage near the reversal potential, which was not the case for this mutant channel. Given the ionic composition of the recording solutions, a possible explanation is that whole-cell calcium currents carried by the L228P mutant may be contaminated by inward cesium permeability currents. This could potentially be further explored by replacing both cesium solutions with NMDG-chloride (internal) and choline-chloride (external).

When the shifts in channel activation and steady-state inactivation are considered independently, they seem to suggest opposing functional consequences. With the shift in activation towards more hyperpolarized potentials, channels would activate at more negative voltages, consistent with a gain-of-function phenotype. On the other hand, the shift in steady-state inactivation towards more hyperpolarized potentials would reduce channel availability, suggesting a loss-of-function phenotype. When the shifts in channel activation and steady-state inactivation are considered together, a complex picture emerges. For example, at a typical neuronal resting membrane potential of -75 mV, most mutant channels would be inactivated, and thus a membrane depolarization would only activate a small fraction of channels. However, in a neuronal circuit comprised of inhibitory and excitatory neurons such as in the thalamus, a GABAergic input would hyperpolarize the membrane, thus triggering massive recovery from inactivation of the channels. Due to the hyperpolarizing shift in half-activation voltage of the mutant channels, a small ensuing depolarization would trigger a massive and selective activation of the mutant channels compared to wild-type, thus potentially leading to rebound bursting, like what is observed with T-type calcium channels and thus hyperexcitability consistent with an overall gain of function (Cain and Snutch, 2013).

It is also important to consider the window current for the L228P mutant, which starts around -70 mV, near the resting membrane potential of neurons, indicating that a fraction of channels at rest are available and open. As such, L228P mutant channels may confer a small yet persistent inward calcium current. Conversely, the window current for wild-type channels begins around -50 mV, and therefore, would largely be inactive at rest or in response to minor subthreshold depolarizations compared to L228P mutant channels. Also noteworthy is that the L228P mutant has a consistent 5% fraction of channels that do not inactivate over a broad voltage range. As a result, the window current of the L228P mutant is much broader than that of wild-type channels, which show complete inactivation. Physiologically, this suggests that L228P mutant channels may contribute to a persistent inward calcium current across a wide range of voltages, even as the membrane potential depolarizes.

Previous research has shown that Cav2.3 channels in CA1 dendritic spines regulate synaptic signalling, in which Cav2.3-mediated calcium influx activates SK channels to dampen synaptic responses (Bloodgood and Sabatini, 2007). Cav2.3-mediated calcium currents have also been shown to increase Kv4.2 currents through increased Kv4.2 channel availability, which attenuates synaptic input and decreases excitability (Murphy et al., 2022). It is plausible that the enhanced subthreshold activity of the L228P mutant Cav2.3 exacerbates these processes, thereby decreasing neuronal excitability. However, it is also plausible that this shift in activation and steady-state inactivation could increase neuronal excitability. Cav2.3 channels are important in forming afterdepolarizations in CA1 neurons of the hippocampus, which are critical for burst firing (Metz et al., 2005). Since L228P mutant channels generally contribute to hyperexcitability, it may be the case that the expression of L228P mutant channels results in increased frequency or strength of afterdepolarizations, which may, in turn, facilitate intrinsic burst firing. Cav2.3 channels have also been shown to mediate the formation of plateau potentials, a transient depolarized state that delays return to rest and have been shown to elicit ictal-type seizure activity (Kuzmiski et al., 2005). It is plausible that the inward calcium current leak of L228P mutant channels stabilizes the plateau potential, increasing the likelihood of ictal-type seizure activity.

When the rate of inactivation for wild-type channels was compared at different EGTA concentrations (5 mM and 0.5 mM, and therefore, differing calcium-buffering capacity), no significant differences were observed, except at +10 mV. This was unexpected, as previous findings have demonstrated that wild-type Cav2.3 channels demonstrate robust CDI (Liang et al., 2003). Comparatively, the wild-type channels in this study exhibited little, if any, CDI. A plausible explanation for the lack of an observable CDI effect in wild-type Cav2.3 channels is that the *CACNA1E* clone used in this study differed from those used by Liang and colleagues.

Specifically, Liang and colleagues used a rat *CACNA1E* construct, whereas this study used a human *CACNA1E* clone (GenBank: NM_001205293.1). Species-specific sequence differences could explain this difference. As observed with the wild-type channel, L228P mutant channels exhibited no CDI.

RFI is an important factor in determining channel availability for certain neuronal firing modes, such as sustained or burst firing, in which wild-type Cav2.3 channels are known to influence burst characteristics (Zaman et al., 2011). However, as discussed earlier, L228P mutant channels contribute very differently from wild-type channels, activating at resting and subthreshold potentials where wild-type channels are inactive. Therefore, the relevance of preserved recovery kinetics needs to be considered in the context of the altered voltage-dependent behaviour of L228P channels and, as such, might not necessarily be relevant to sustained or burst firing.

The L228P substitution clearly has profound effect on the voltage dependence activation, inactivation, and potentially on permeation. Residue 228 is located towards the extracellular end of the domain I S5 segment, a region that contributes to the formation of the channel’s pore (Zamponi et al., 2015). This may result in altered ion selectivity for calcium over monovalent ions such as cesium. While not localized within the voltage-sensing domain of the channel, the insertion of the proline residue would be expected to cause significant structural alterations that may lead to facilitated movement of the domain I voltage sensor, thus perhaps accounting for the enhancement of channel activation. Inactivation of the channels is thought to involve interactions between the domain I-II linker and S6 segments of the channel (Zhang et al., 1994; Stotz and Zamponi, 2001). It is possible that allosteric coupling between residue 228 and the inactivation machinery of the channels may occur. Alternatively, the change in voltage-dependence of inactivation may be secondarily due to strong coupling between activation and inactivation.

Taken together, our findings demonstrate that the L228P mutation alters channel activity and availability across a broad range of voltages. These findings contribute to ongoing studies of Cav2.3 channel mutations and their impact on channel biophysics. Ultimately, continued study of Cav2.3 channel mutations is needed to aid the development of precision therapies for *CACNA1E*-associated DEE conditions.

## Author contributions

GWZ, IAS and DK conceptualized the study. DK carried out experiments, performed statistics and analyzed data. MG and LF assisted with data analysis. DK and GWZ drafted the manuscript. DK, IAS, MG, LF and GWZ edited the manuscript. GWZ secured funding and supervised the study. All authors read and approved the final manuscript.

## CRediT authorship contribution statement

Devon Khousakoun: original draft, methodology, investigation, formal analysis, data curation, conceptualization.

Ivana A. Souza: conceptualization, writing-review and editing

Maria A. Gandini and Laurent Ferron: Methodology, validation, writing-review and editing

Gerald W. Zamponi: Writing-review and editing, supervision, project administration, funding acquisition.

## Funding

This work was supported by a grant to GWZ from the Canadian Institutes of Health Research (CIHR). DK holds a CIHR studentship.

## Declaration of competing interest

The authors declare that they have no known competing financial interests or personal relationships that could have appeared to influence the work reported in this paper.

## Data availability

The authors confirm that the data supporting the findings of this study are available within the article

## Notes

### Competing Interest Statement

The authors have declared no competing interest.

